# Temporal evolution of Neural Codes: The Added Value of a Geometric Approach to Linear Coefficients

**DOI:** 10.1101/2025.10.10.681582

**Authors:** Théo Desbordes, Itsaso Olasagasti, Nicolas Piron, Sophie Schwartz, Nina Kazanina

## Abstract

Multivariate decoding analyses have become a cornerstone method in cognitive neuroscience. When applied to time-resolved brain imaging signals, they provide insights into the temporal dynamics of information processing in the brain. In particular, the temporal generalization (TG) method—where a decoder trained at one time point is tested on others—is commonly used to assess the stability of neural representations over time. However, TG performance can be ambiguous: distinct representational dynamics—such as sparse versus distributed activity, or scaling of activity versus recruitment of new units—can yield similar TG matrices. Moreover, even when generalization is strong, underlying neural representations may still be evolving in ways that TG alone fails to reveal. This ambiguity of performance profiles can mask meaningful changes in the geometry of neural representations. In this study, we use controlled simulations to demonstrate how different dynamic processes can produce indistinguishable TG profiles. To resolve these ambiguities, we propose a complementary approach based on the geometry of the learned linear coefficients. Specifically, we quantify the **Rotation Angle θ** between decision subspaces (with cosine similarity) and the **Feature Density α** (capturing whether feature contributions are distributed or sparse). Together, these measures complement TG analyses, revealing how neural representations evolve in space and time. Beyond time-resolved decoding, our approach applies broadly to any linear model, offering a geometric perspective on representational dynamics.

**Graphical abstract:** 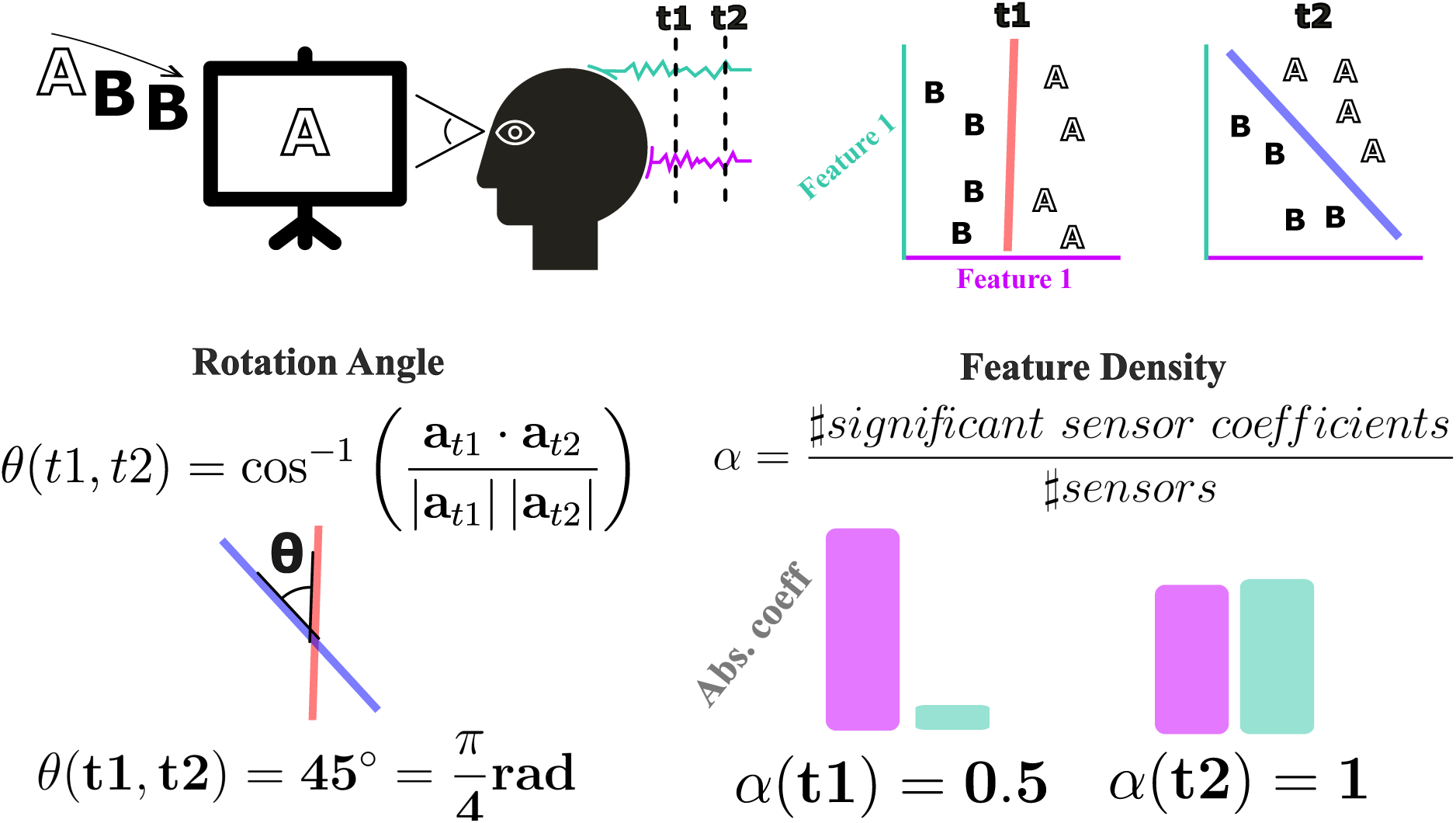

## 1. Introduction

**Neural representation** is a central concept in contemporary neuroscience (Hubel & Wiesel, 1962; Marr, 1982; deCharms & Zador, 2000; Kragel et al., 2018). It refers to patterns of neural activity that encode external and internal states, providing the basis for the brain’s processes that support interpretation, prediction, and interaction with the world. How these internal representations are formed has intrigued scholars since antiquity; Aristotle, for example, likened the mind to wax receiving the impression of a signet ring.

Behaviorism emerged in the early 20th century as a dominant approach in psychology, emphasizing the study of observable behaviors over internal mental states, which were seen as unreliable and irrelevant. However, since the late 1950s and increasingly with the advent of neuroimaging in recent decades, research has shifted toward investigating neural representations. This modern focus acknowledges that behavior is deeply rooted in brain processes, allowing researchers to bridge observable actions with the mental and neural mechanisms that produce them. While this notion underpins much of modern cognitive neuroscience, key questions remain regarding how best to quantify and track internal codes over time.

Yet, the term representation may refer to various mental and/or neural processes (Favela & Machery, 2023, 2025). Here we will adopt a working definition of representations as patterns of neural activity that carry information about external or internal states, potentially useful for the brain to guide perception and behavior, consistent with previous usage (Vilarroya, 2017; Heinen et al., 2024).

Encoding and decoding models of brain activity are major instruments in the neuroscientist’s toolbox to probe for neural representations (Rolls & Treves, 2011; Tong & Pratte, 2012; Mathis et al., 2024). **Encoding models** aim at predicting neural activity from experimental parameters, and as such they are also called **forward models** (from stimulus to neural activity). **Decoding models** do the reverse: they attempt to recover experimental variables from the corresponding recorded neural activity, and thus are called **backward models** (from neural activity back to the stimulus). Formally, these models are often rooted in the same statistical modeling framework. Classical examples include regularized linear and logistic regressions. Both types of models aim to predict a **target** as a weighted sum of **features**, with the **weights** learned directly from the data. However, they differ in what is used as features and targets. In decoding or backward models, features can be any kind of direct or indirect neuronal measurement: single-neuron firing rates, local field potentials, voxels from functional Magnetic Resonance Imaging (fMRI), etc. The target of decoding models (what they aim to predict) are experimental variables of interest to the researcher. In the simplest case, it can be classifying two categories of trials (e.g., “present” vs “absent” stimulus, “correct” vs “incorrect” behavioral performance, …). **Figure 1A** shows an example with two features and a target consisting of two classes.

**Figure 1:**
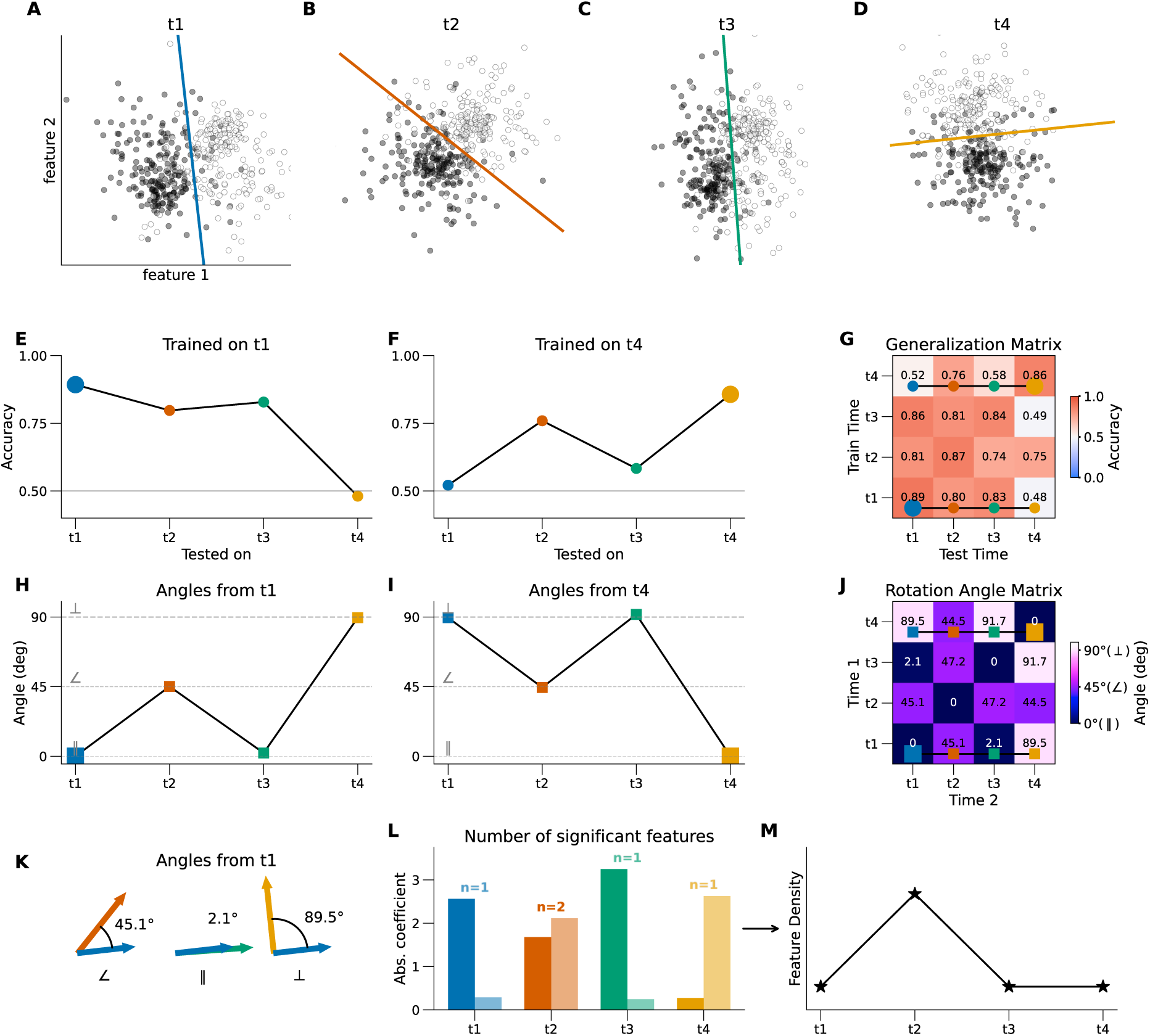
Didactic example illustrating decision boundaries, generalization performance, Rotation Angle and Feature Density. ***A-D****: Illustrative data from four time points and their corresponding decision boundaries, i.e., when a logistic regression classifier is trained to differentiate two experimental conditions using recorded data at each of these time points, separately. Empty and filled round markers correspond to data points from the two classes. The data is 2-dimensional, in this toy example the model has two input features, corresponding to two sensors or source estimates. The decision boundary for each time is shown with a thick solid line*. ***E-F****: classification accuracy for the decoders trained at t1 (E) and t4 (F) and evaluated on data from all time points. These correspond to the bottom and top lines of the generalization matrix in G, respectively*. ***G****: Temporal Generalization matrix, showing the decoding performance of each classifier trained at one time (y-axis) and tested at each time (x-axis)*. ***H-I****: angle between the decision boundary at t1 and that of each other time point (H), and between t4 and each other time point (I). The plotted values correspond to the bottom and top rows of the angle matrix in J, respectively. Note that the angle between the same coefficient vector (same time) is equal to 0, by design*. ***J****: Rotation Angle matrix, showing the angle between each pair of time points. Note that the diagonal is necessarily filled with zeros (angle between a coefficient vector and itself). Unlike to the generalization matrix, the angle matrix is symmetrical, by design*. ***K****: Illustration of the Rotation Angle calculation. The angle is based on the cosine similarity between the coefficient vector from t1 (blue), and that of each other time point*. ***L****: Absolute value of the coefficient associated with feature 1 (darker shade) and 2 (lighter shade) for each time point. On top of the bars for each time points is written the number of coefficients with magnitudes that significantly differ from 0*. ***M****: Feature Density for each time point, defined as fraction of features (relative to the total number of features) with coefficients significantly different from zero*.

Brain decoding may also refer to a related field of research, which has a quite different aim and distinct applications, such as reconstructing natural perceptions (Défossez et al., 2023; d’Ascoli et al., 2024; Lévy et al., 2025) and rehabilitation of impaired brain function (Anumanchipalli et al., 2019; Moses et al., 2021; Lorach et al., 2023). The methods used for this type of decoding are usually deep learning models that are highly expressive, at the cost of reduced interpretability. This is in stark contrast with the simple, often linear methods used when brain decoding is applied to infer neural representations as in the present paper.

Time-resolved decoding using non-invasive methods such as electroencephalography (EEG) and magnetoencephalography (MEG) (Baillet, 2017; Gross, 2019), has been particularly successful in dissecting fast sequences of neural events (Gwilliams & King, 2020; Honari-Jahromi et al., 2021; Desbordes et al., 2024). Time-resolved decoding consists in fitting statistical learning models (typically classification or regression linear models, thereafter called “**decoders**”) at each timestep of these millisecond-resolved signals. Above-chance decoding performance at a given time point indicates that information related to a specific experimental condition is present in the neural signal at that moment. Hereafter, we will take the view that significant decoding performance offers primary evidence for the presence of a neural **representation.** While decoding performance is a necessary condition for the identification of a representation, we should keep in mind that it is not sufficient, as additional criteria—for instance, causal relevance—must also be met (Ritchie et al., 2019).

The **Temporal Generalization (TG)** method (King & Dehaene, 2014) is an extension of time-resolved decoding that allows testing for the temporal stability of representations. The underlying assumption is that if a classifier trained at one time point successfully generalizes to another time point, this is evidence for the information being encoded in a similar way at these two time points, i.e., a “stable representation” (as shown in **Figure 1E-G**, t1, 2, t3). Conversely, if a classifier trained at one time does not generalize to another time point then this is evidence for the information being encoded differently at these two time points. In other words, if classifiers trained at different times both have good **within-time performance** (trained and tested at the same time), but chance-level **across-time performance** (tested at each other’s time), then this shows that information is present but encoded differently at each time point (**Figure 1E-G**, t1, t4). TG have been widely used in time-resolved imaging (Marti et al., 2015; Rosenthal et al., 2020; Gwilliams et al., 2022; Bo et al., 2022) and have been argued to be particularly useful to investigate processes with rapid cascades of neural events, such as language understanding (Fyshe, 2020).

Put simply, time-resolved (within-time) decoding tells **when** information is present, but not **how** representations evolve. TG (across-time decoding) addresses this limitation by probing the **stability** of representations over time, yet it remains agnostic to the underlying feature dynamics. As we will see, failures of generalization can arise for multiple reasons, while robust generalization may mask changes in the coding scheme.

Thus, while within-time performance and TG provide valuable insights into the timing and stability of decodable information, they fall short of capturing how representational structures themselves transform or persist. By reducing comparisons to a single performance metric, these approaches discard the rich geometric information embedded in the linear decoders. We argue that examining this geometry can offer crucial complementary insights into the evolution of neural representations. In what follows, we describe how such geometric information can be recovered from the learned coefficients.

A linear decoder assigns a scalar coefficient (weight) to each input feature (e.g., each MEG sensor or source estimate), specifying how they are combined to best predict the target from the available data. This coefficient vector of a classifier defines a **decision boundary** in feature space (more precisely, a **hyperplane**). In **Figure 1A-D**, we illustrate this process with synthetic data consisting of two features and four time points and their corresponding decision boundary. Classical TG consists in applying a classifier trained at one time point to the data at another time point, effectively assessing which side of the previously established decision boundary new data points fall onto, and computing the classification performance (**Figure 1E and 1F** show the generalization of decoders trained at t1 and t4, while Figure **G** shows the full TG matrix). Our methodological extension is to analyze the decision subspaces at the two time points directly through two geometric measures: (i) their relative **Rotation Angle** (measured with cosine similarity) and (ii) the **Feature Density** (the proportion of coefficients with significant magnitude).

In practice, the Rotation Angle **θ** is measured using cosine similarity between pairs of activation pattern vectors (**Figure 1K**; see Methods). This measure is naturally correlated with generalization performance: orthogonal decision boundaries often yield chance-level generalization, and perfectly aligned hyperplanes often mean good generalization. However, the correlation is not perfect. For example, in **Figure 1H** the angle between the first and second time points is equal to 45 degrees, while the one between first and third time points is close to zero. This is easily visible when looking at the decision boundaries: t1 (**Figure 1A**) and t3 (**Figure 1C**) are aligned, whereas t2 (**Figure 1B**) is tilted compared to the other two. Despite this substantial geometrical difference generalization performance is similar (0.80 and 0.83 accuracy for t2 and t3; **Figure 1E and 1G**).

In the same simulation, the angle of the fourth time point is orthogonal to the first and third time points (**Figure 1I**). However, the generalization performance from t4 to t1 is basically at chance (0.52 accuracy), whereas that from t4 to t3 is above-chance (0.58 accuracy). These two examples illustrate that the performance of decoders and the angle between their decision hyperplanes can differ substantially, in a way that could be overlooked if only computing the decoding accuracy. In other words, the relationship between the geometry of the decision boundaries and generalization performance is not straightforward.

Another key yet often overlooked attribute of linear models’ coefficients is how **sparse** they are. Do all features (sensors or sources) contribute equally, or are the weights concentrated in just a few? To quantify this, we introduce the **Feature Density α** (**Figure 1L and M**, see Methods). This measure gives a numerical estimate of how distributed or sparse a decision subspace is. In the simplified example shown in **Figure 1**, this corresponded to estimating if a single feature contributes to the decision, or whether the two features are necessary. Thus, unlike the Rotation Angle measure, which applies to a pair of time points, the Feature Density is a property defined at each individual time point. In our example data, both features are critical for the classifier only at t2 (**Figure 1L** and M), whereas the other time points largely rely on a single feature (characterized by basically horizontal or vertical decision boundaries on **Figure 1A, C and D**).

The contributions of the paper are:

◦ Illustrate potential ambiguities associated with performance metrics of linear models and the standard generalization methods, focusing on TG, by providing simulated examples
◦ Investigate the geometry of the decision subspaces to disambiguate these cases, namely compute the **Rotation Angle** between pairs of pairs of activation pattern vectors (measured with cosine similarity) and **Feature Density** (proportion of features significantly contributing to the decoder).
◦ Suggest use cases for this approach beyond time-resolved decoding.

Overall, we demonstrate that characterizing the decision subspace geometry provides critical insights into the temporal evolution of neural codes, providing an additional layer of interpretation for time-resolved decoding models.

## 2. Methods

### 2.1. Simulation Framework

We simulated multivariate brain data to model temporal evolution of neural activity patterns. Simulated datasets were generated with the following parameters: **300 sensors**, **100 trials of each of the two classes**, and **100 time points** per dataset (1 s at 100 Hz). Patterns were introduced during the **active time window**, defined as time points **between 0.1 s and 0.9 s**. This corresponds to a realistic amount of data for a typical M/EEG experiment.

At each time point within the active time window, 10% of the sensors (30) were assigned nonzero activity patterns associated with the experimental condition for all conditions, except *Distributed Cascade*, where one third of the sensors (100) were affected instead (see below). The base patterns were sampled from a standard normal distribution and evolved following one of six predefined scenarios:

◦ **Sparse Cascade (Figure 2A)**: each of the sensors contributing to the pattern is active only for a short duration (corresponding to the length of the active time window divided by the number of sensors in the pattern), with an overlap of two timesteps. This mimics a chain of sparse, independent processes, e.g. neurons firing sequentially (Abeles, 1982; Diesmann et al., 1999).
◦ **Distributed Cascade (Figure 2B**): a new random pattern is sampled at each time point, affecting a third of all sensors. This high-dimensional random assignment promotes orthogonality of the signal across consecutive time points. This is akin to the reservoir computing framework (Maass et al., 2002; Jaeger & Haas, 2004), where neural populations produce high-dimensional, quickly changing activity patterns.
◦ **Sustained (Figure 2C**): the same sensors stay active with constant amplitude throughout the active time window. This is akin to persistent activity in working memory (Goldman-Rakic, 1995; Constantinidis et al., 2018).
◦ **Progressive Switch (Figure 2D**): an initial “early” pattern starts fully active and progressively fades out, while a second “late” pattern starts weakly active and grows in strength. The “early” pattern is scaled down by a factor that linearly evolves in time from 1 (beginning of the active time window) to 0.2 (end), while the “late” pattern starts weakly active and is scaled up by a factor that linearly evolves in time from 0.2 to 1. This is akin to a transfer of information from one subspace to another, or progressive change of representation, e.g. an initial “sensory” representation is gradually replaced by a “memory” representation (Xie et al., 2022; Chen et al., 2024).
◦ **Scaling (Figure 2E**): the pattern’s amplitude linearly increases from zero to one across the active time window. In other words, the direction of the signal stays the same, but its amplitude grows over time. This is akin to ramping activity often observed in single-units in the planning and control of motor movements (Narayanan, 2016; Affan et al., 2025), and has also been hypothesized to represent external variables in entorhinal cortex (Nolan, 2025)
◦ **Recruitment (Figure 2F**): sensors are linearly recruited into the pattern throughout the time window, with more initially inactive sensors becoming progressively involved. This is akin to the gradual recruitment of new units during semantic integration (Desbordes et al., 2023).

**Figure 2:**
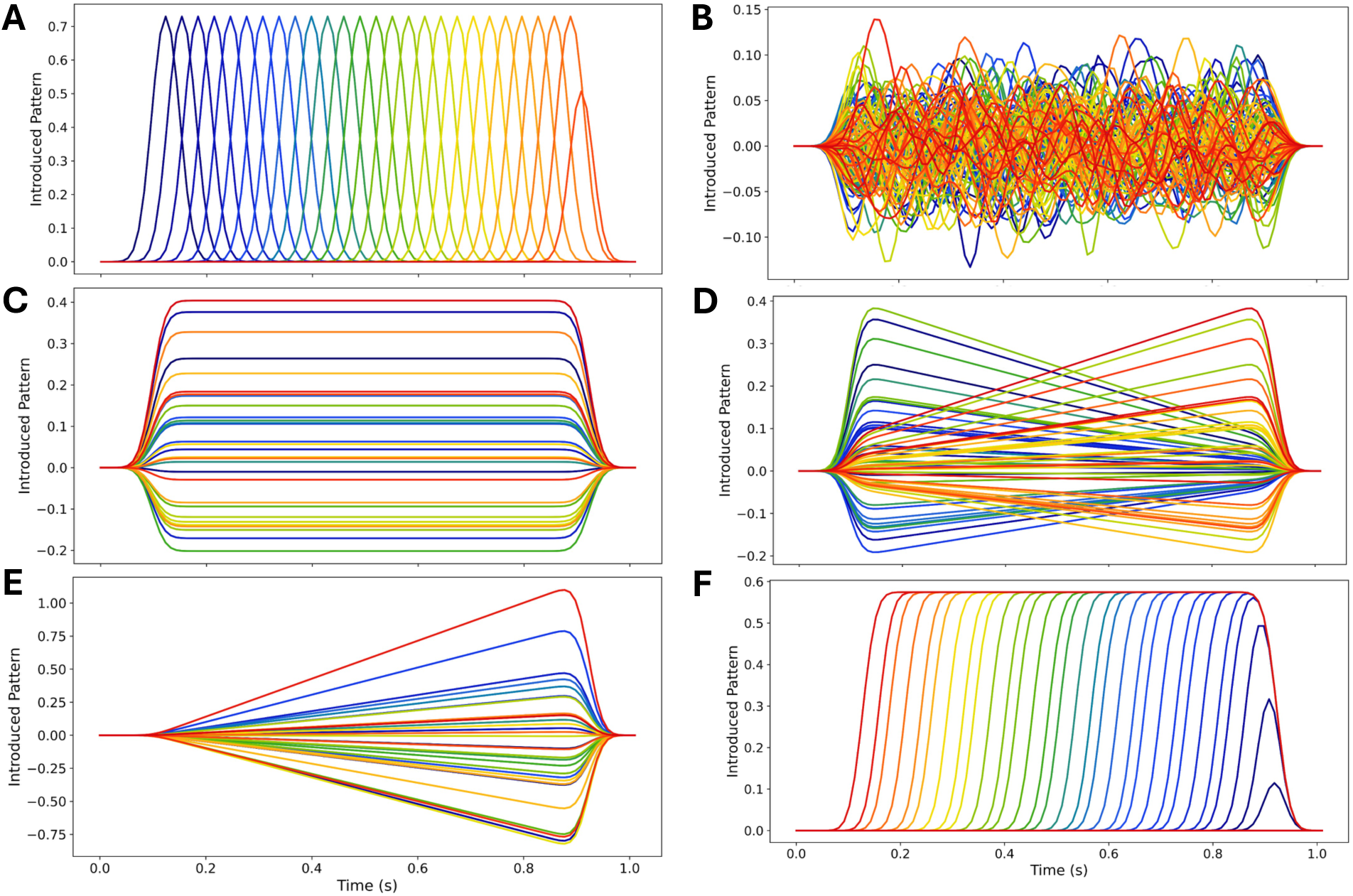
Patterns introduced in each simulated scenario. *Differential of activity between the two conditions in each trial, before the addition of noise, for every scenario*. ***A****: Sparse Cascade, **B**: Distributed Cascade, **C**: Sustained, **D**: Progressive Switch, **E**: Scaling, **F**: Recruitment*.

Each simulated dataset included subject-specific and trial-specific normally distributed noise. To approximate the smooth temporal autocorrelation typically observed in M/EEG data, the resulting data matrices were temporally smoothed using a Gaussian kernel with radius σ = 2. This was the final step of the data generation procedure, applied after the introduction of the patterns and the noise. Trial average activity for each scenario can be found in Supplementary Figure 10.

### 2.2. Classifier Training and Temporal Generalization

At every time point, we trained separate L1- and L2-regularized Ordinary Least Squares classifiers, Logistic Regressions, as well as linear Support Vector classifiers (using scikit-learn’s implementation) to discriminate between two conditions. In the main text we present the results for the L2-regularized Logistic Regression. Results were robust across model initializations and classifier types (see Supplementary Figure 4-10). Note that in some neuroimaging decoding studies, the normalization prior to classifier training, and/or the optimization of hyper-parameters (like the regularization strength), is done separately for each time point. Here, to ensure comparability across time, normalization and hyperparameter tuning were held constant across all time points..

These trained classifiers were then evaluated at all other time points, yielding a **temporal generalization matrix** (**Figure 1G**) that reflects decoding performance over time (King & Dehaene, 2014).

Classifier weights are difficult to interpret directly because correlated noise can produce large coefficients on sensors that contrain only noise (Hebart & Baker, 2018; Kriegeskorte & Douglas, 2019). We therefore applied the Haufe et al. (2014) transformation, multiplying weight vectors by the data covariance matrix to obtain **activation patterns** — interpretable forward mappings of signal contributions. The analysis of activation patterns has been used in numerous studies to identify the spatial localization of information encoding in the brain (Čeko et al., 2022; Wen et al., 2024). The resulting patterns are interpretable, in the sense that a feature associated with a high value contains information about the condition of interest and participates strongly in the decoder’s prediction.

Formally, to interpret classifier weight vectors **w**, we multiplied them by the empirical data covariance matrix Σ_!_ to recover **activation patterns** (following Haufe et al., 2014).

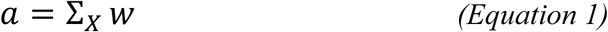

These transformed weight patterns, subsequently called the models’ **coefficients**, were used in all subsequent analyses. The vector of coefficients corresponds to a geometrical object that we call the **decision subspace** (as opposed to decision boundary, before the transformation). Like the feature normalization, the data covariance was estimated jointly from all time points to enable comparison among the resulting coefficients. We show recovered activation patterns for each scenario in **Supplementary Figure 1**.

### 2.3. Contributing Sensors Selection

We identified sensors contributing significantly to decoding at each time point by comparing active versus baseline periods. Significance was assessed with one-sample t-tests comparing the average absolute classifier coefficients during the pattern-active period to those during the baseline period (pre-onset, 0 s to 0.1 s). Significant sensors (p<0.01 with False Discovery Rate (FDR) correction for multiple comparisons (Benjamini & Hochberg, 1995)) were selected for further analyses. This step prioritizes the decision subspace, rather than the random (noise-related) coefficient fluctuations that would dilute subsequent geometrical measures.

### 2.4. Quantifying Rotation Angle and Feature Density

To characterize the temporal evolution of the patterns, we computed:

◦ **Rotation Angle** (**θ)**, based on the cosine similarity between the coefficient vectors at two distinct time points, in radians:

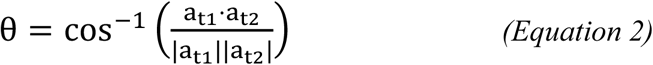

An angle **θ = 0** indicates perfectly **aligned** patterns, θ = 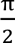 *rad* = 90° indicates **orthogonal** patterns, and θ = π *rad* = 180° indicates perfectly **reversed** patterns. For random unrelated patterns θ ≈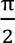 *rad* = 90° is expected. A full representation of angular differences between pairs of time points yields a symmetric matrix of the same size as TG, with zeros on the diagonal (the activation pattern is always aligned with itself), and off-diagonal values indicating alignment of patterns trained at different times. We call it the **Rotation Angle matrix**.

To characterize the proportion of features (here, sensors) contributing significantly to the decoding at each time point, we computed:

◦ **Feature Density** (**α)**: proportion of features with coefficient whose magnitude are significantly different from zero. In regularized linear models, we expect features containing noise but no signal to have a coefficient close to zero. We performed one-sample t-test comparing the absolute classifier coefficients against zero, separately for each time point (p<0.01 with False Discovery Rate correction). To reduce the number of comparisons, we only tested the sensors selected by the Contributing Sensor Selection. The ratio of the number of significant sensors to the total number of sensors gives an estimate of how **distributed** (many sensors carrying the signal) or **sparse** (few sensors carrying the signal) the pattern is.

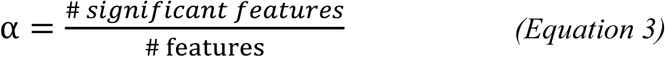 Where a high value of α means distributed contribution (α = 1 implies that all features contribute), and low values are associated with sparse coefficients (α = 0 means that no feature contain signal). This measure is computed independently at each time point, yielding a time course of feature density over time that may be used to further characterize the neural dynamics (see Results section). This measure is highly sensitive to the regularization of the linear model—stronger regularization produces sparser solutions. Therefore, it is essential to estimate the regularization hyperparameter across all time points and apply the same value consistently.

All the code for the simulations, analyses, and plotting will be made freely accessible upon publication. They consist of simple-to-use, flexible Python functions based on NumPy (Harris et al., 2020), SciPy (Virtanen et al., 2020), and MNE-Python (Gramfort et al., 2013), and a jupyter notebook tutorial.

## 3. Results

### 3.1. Sparse vs. distributed Cascade Scenarios

We first compared two dynamic scenarios with rapidly changing activity patterns: the **Sparse** and **Distributed Cascades**. In the **Sparse Cascade**, different sensors represent the current class at different times, mimicking sequential information transfer. (**Figure 2A**). At each time point, a distinct sensor carries the signal, with a small overlap for neighboring time points due to the temporal smoothing. In the **Distributed Cascade** however, the same subset of sensors is active over the whole time window, but with quickly changing activity: a new random pattern differentiates the two classes at every time point, akin to chaotic activity in the same neural population (**Figure 2B**).

Each scenario produced comparable within-time decoding performance (**Figure 3A**). Both yielded similar Temporal Generalization (TG) profiles: high accuracy along the diagonal but poor off-diagonal generalization (**Figure 3B & 3C**), consistent with quickly changing neural patterns associated with orthogonal decision subspaces (**Supplementary Figure 2**). By design, there was a difference in the number of sensors involved in each scenario, and that was well captured by the Feature Density (**Figure 3D**), i.e., one sensor at a time for the sparse cascade and closer to a third of the sensors at each time point for the distributed cascade. Note that at any given time, there are several sensors that do not significantly contribute to the pattern (contribution close to zero in **Figure 2B**), which explains that we don’t reach α = 1/3.

**Figure 3:**
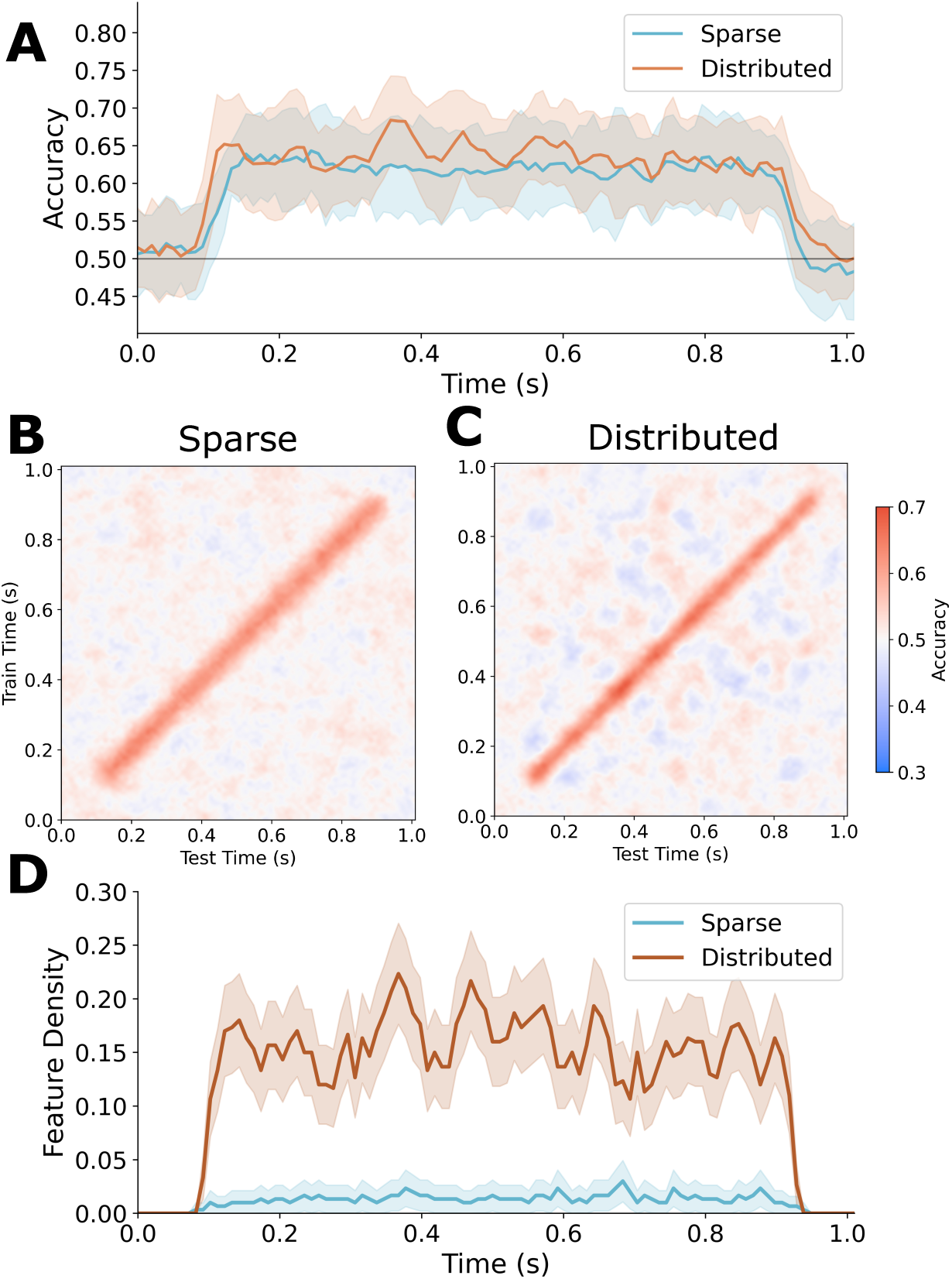
Classifier performance, generalization and Feature Density for Sparse and Distributed Sequential scenarios. ***A****: Within-time decoding performance for both scenarios. Temporal generalization matrices for Sparse Cascade (**B**)and Distributed Cascade (**C**). **D**: Feature Density over time showing highly distinct indexes of the sparsity of the decision subspace for the proposed scenarios*.

Direct comparison of introduced (**Figure 2A & 2B**) and recovered patterns (**Supplementary Figure 1A & 1B**) confirmed accurate pattern recovery.

To sum up, using the Feature Density, an estimate of the number of sensors significantly contributing to the classifiers’ decision, we successfully distinguished between a sparse vs a distributed sensor scenario, while they both elicited similar TG maps.

### 3.2. Sustained versus Progressive Switch Scenarios

Next, to illustrate that a stable TG does not always mean perfectly stable or recurrent underlying dynamics, we compared the case of a stable pattern (**Sustained, Figure 2C**) with that of a transmission of information from one subspace to another (**Progressive Switch, Figure 2D**). The latter case can be seen as an example of the transmission of information from sensory to associative regions, whereby an “early” sensory pattern would first appear, before it is progressively replaced by a “late” memory pattern carried by a completely distinct set of sensors.

Both the **Sustained** and **Progressive Switch** scenarios exhibit a good within-time decoding performance (**Supplementary Figure 3**) and a square generalization matrix (**Figure 4A-B**). A careful inspection showed that, after the transformation has been completed (t = 0.9 s), the generalization performance of a decoder trained at the beginning of the active window is slightly weaker for the Progressive Switch case, illustrating the change of strength of the two patterns. But the performance is still significantly better than chance. This is anticipated: the “early” pattern is still present (albeit about five times weaker than at the beginning of the active time window), and in a good Signal-to-Noise condition, above chance performance is to be expected. This highlights that the within-time and generalization performance metrics may overlook drastic differences in underlying pattern dynamics.

**Figure 4:**
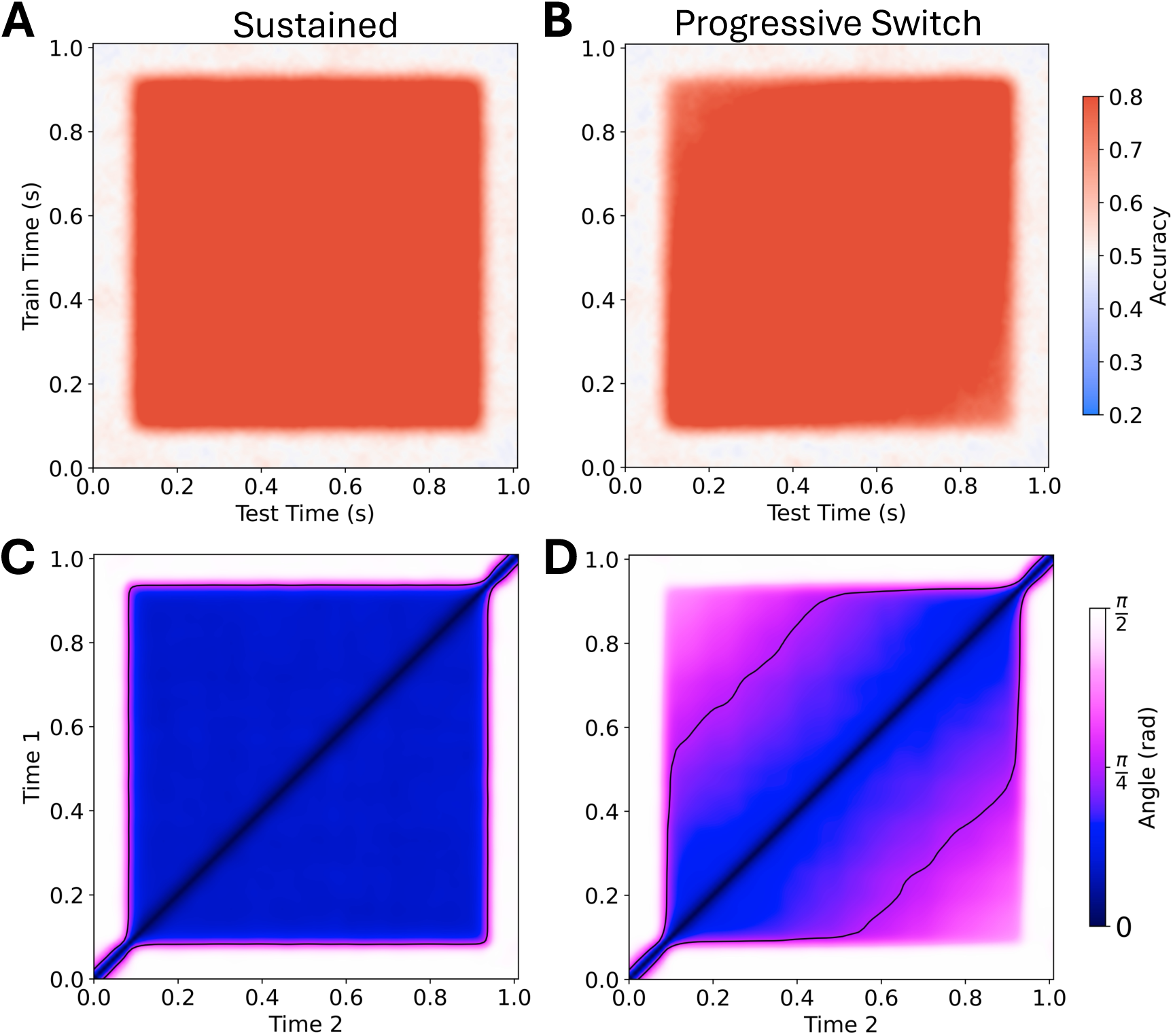
Decoding generalization performance and decision subspace Rotation Angle for Sustained and Progressive Switch scenarios. *Temporal generalization matrices for Sustained (**A**)and Progressive Switch (**B**) scenarios. Rotation Angle matrices for Sustained (**C**) and Progressive Switch (**D**). The thin black lines represent isoline of θ* = 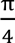 *rad* = 45°. *Note that although the temporal generalization profiles are similar in these two scenarios, the Rotation angle differs, correctly characterising the progressive transfer of representation in the Progressive Switch*.

Looking at the Rotation Angle matrices resolves the ambiguity. For the Sustained scenario (**Figure 4C**), we see no change in orientation: the angle between the decision subspaces at any pair of time points during the active window is close to zero. This illustrates that the subspace does not change over time. By contrast, for the Progressive Switch, the angle progressively changes as the second pattern is introduced, with orthogonal angles after the full transformation (between t = 0.1 s and t = 0.9 s) and perfect alignment on the diagonal, and a smooth transition in between. For example, at the halfway point of the scenarios (t = 0.5 s), we have an angle close to 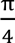 *rad* (45°) with both t = 0.1 s and t = 0.9 s. Thus, the continuous transfer of class information from the “early” to the “late” subspace is revealed with the measure of the Rotation Angle between pair of decision subspaces.

### 3.3. Scaling vs. Recruitment Scenarios

Finally, we contrasted two ramping-like dynamics, or an increasingly stronger square generalization matrix. This TG profile has been identified in previous studies and interpreted as integration of semantic information through the recruitment of new units in a compositional cell assembly (Desbordes et al., 2023). Moreover, single-unit or single sensor signatures compatible with a ramping pattern (progressive increase of signal) are seen in various domains e.g. accumulation of evidence in decision making (Gold & Shadlen, 2007; van Vugt et al., 2012; Brody & Hanks, 2016). How can we differentiate cases where ramping neural signals come from increased activity in single sensors, and cases where it is due to the recruitment of an increasing number of sensors?

We describe the case of **Scaling** of a small number of sensors, starting with low values and progressively increasing to full strength (**Figure 2E**), and the case of **Recruitment**, where sensors are progressively added to the representation and remain active until the end of the time window (**Figure 2F**). These are two examples of progressive divergence in the neural representations, but with different underlying sources. As such, they are associated with similar TG matrices (**Figure 5A-B**).

**Figure 5:**
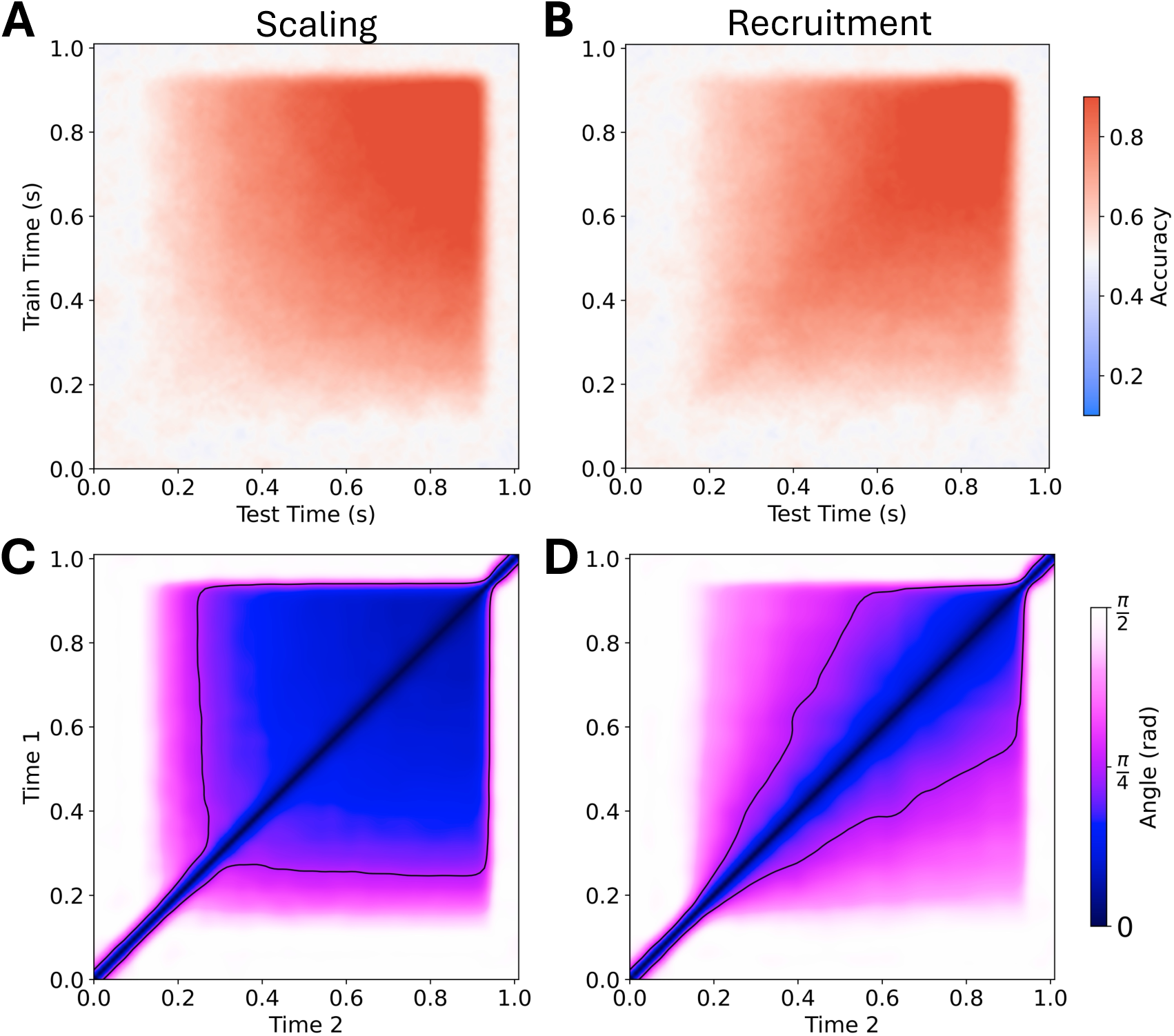
Classifier performance and decision subspace Rotation Angle for Scaling and Recruitment scenarios. *Temporal generalization matrices for Scaling (**A**)and Recruitment (**B**). Rotation Angle matrices for Scaling (**C**) and Recruitment (**D**). The thin black lines represent isoline of θ* = 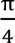 *rad* = 45°*. Note that for Scaling (C), after t = 0.3 s there is hardly any change in orientation of the decision subspaces (low angles), whereas in the case of Recruitment (D) for any row, there is an evolution of the color witnessing the change in the angle*.

In the case of Scaling, the signal direction present in the affected channels is fixed, only the magnitude grows linearly. For this we predict that the angle should be close to zero across the entire time window. Indeed, as soon as there is enough signal (at t > 0.3 s), the angles have values below 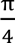(45°), consistent with a stronger alignment of the decision subspaces (**Figure 5C**). In the case of Recruitment, the addition of new sensors has a stronger impact on the changes in orientation of the subspaces over time, especially early on when few sensors are contributing (**Figure2D)**.

Across these three pairs of examples, we showed that new geometrical measures (Feature Density and Rotation Angle) computed on the decision subspaces can clearly distinguish between cases with similar generalization performance, but with distinct underlying dynamics of representations. Owing to its links with meaningful neural dynamics, this geometrical framework provides unprecedented support for the interpretation of complex temporal patterns of neural activity.

## 4. Discussion

In this study, we emphasized important but often overlooked ambiguities associated with the widely used temporal generalization (TG) method in neural decoding analyses. Through simulations of non-invasive brain recordings, we demonstrated that multiple scenarios of canonical neuronal processes can produce comparable TG profiles, blind to the underlying nature of information encoding. To address this limitation, we proposed complementary analyses based on the geometry of the classifiers’ decision subspaces. **Feature Density** can set apart scenarios where the information is encoded in a sparse or distributed fashion (**Figure 3**). **Rotation Angle** can tease apart cases where apparent stability in TG obscures an evolving representation (**Figure 4**), as well as cases where different coding schemes give similar TG signatures (**Figure 5**). These results highlight that decoding performance is necessary but not sufficient for understanding neural dynamics: geometry provides a missing link.

Our approach adds a geometric perspective to time-resolved decoding. By focusing on the orientation and sparsity of learned subspaces, it connects decoding analyses with recent theoretical work on neural population geometry (Chung & Abbott, 2021). Although developed for temporal decoding, the same decomposition can be applied to any collection of linear mappings. In general, we contend that this method will be useful in cases where the data is high-dimensional and/or where each feature does not have a clear associated meaning, making the interpretation of individual coefficient challenging. This is true for multivariate brain decoding, but also for many other classic analyses in neuroscience and beyond.

Example applications beyond multivariate brain decoding include: projection from raw sensory inputs onto a low-dimensional latent space (e.g. via principal component analysis; Jolliffe, 2011), the alignment of activity patterns across two brain regions using canonical correlation analysis (Thompson, 2000), the mapping between successive time points in a neural population under the assumption of linear dynamics (linear state-space models; Smith & Brown, 2003), and metric-learning transformations (Lakretz et al., 2018; Jalouzot et al., 2024; Salle et al., 2024). This approach could also be extended to encoding analyses, particularly in contexts where the input features are high-dimensional — such as word embeddings from large language models — and direct interpretation is challenging. One good use case could be the growing trend in using high-dimensional embeddings from Artificial Neural Networks (ANNs) to predict brain activity (Huth et al., 2016; Schrimpf et al., 2020; Zhuang et al., 2021; Caucheteux & King, 2022; d’Ascoli et al., 2025). Moving beyond neuroimaging, decoding approaches have also recently emerged in ANNs to analyse their inner workings, generally called “probing” analyses. For example, many linguistic features were identified as linearly decodable in Large Language Models (Lakretz et al., 2019; Müller-Eberstein et al., 2022). Here as well, looking at the geometry of the representation could give further insight into how these features are represented in ANNs. For example, it could be applied to track representational drift across layers or time in recurrent models. Another approach consists in “stitching” two models at a certain layer with a linear mapping, and our method could allow investigating the geometry of such transformation (Lenc & Vedaldi, 2015; Bansal et al., 2021)

A current limitation is that our method applies strictly to linear models, where the decision subspace is well-defined. Extending geometric analyses to deep networks, where each layer implements a sequence of linear transformations followed by nonlinearities, remains an open challenge. Nonetheless, investigating the evolution of such transformations could yield valuable insights into the computations performed by hierarchical models.

It is worth noting that in neuroimaging studies, non-perfectly square TG patterns are often taken as signs of stability (Oh et al., 2019; Weaver et al., 2019). Here, we showed that even when the TG closely resembles a perfect square, there might be major changes in the underlying representation (**Figure 4B & D**)

Time-resolved decoding has been extended in various ways, including temporally-delayed linear models (Liu et al., 2021), that aim to identify sequential reactivations of states, and temporal pattern classification and generalization across locations (Santos-Mayo et al., 2025), that look for class-related information in short temporal sequences. These methods serve different goals and would not help distinguish the scenarios considered in the present study. Also related to the proposed method, is the Representational Similarity Analysis (Kriegeskorte et al., 2008; Kriegeskorte & Kievit, 2013), that also looks at the geometry of representations through measures of similarity based on second-order isomorphisms (Shepard & Chipman, 1970).

Regarding our choice to use a method based on cosine similarity for the estimation of angles, we selected this relatively simple method because it yields similar outputs when compared to conceptually more complex methods such as Direct Projection, and estimation of a linear transformation between pairs of coefficients followed by Polar Decomposition or Procrustes (Gower, 1975). However, note that in the case of high-dimensional spaces, multiple principal angles would provide a more thorough description (than a single angle approach). Methods like Procrustes can also isolate the rotation matrix from the linear transformation between two hyperplanes, giving a much richer view on the rotational components between the two subspaces. In the simple cases considered here, looking at a single, “aggregate” angle was sufficient, but it is likely that advanced geometrical descriptions, that are beyond the scope of this paper, will necessitate to consider this aspect. Similarly, an alternative measure of sparsity used in economics is the Gini coefficient (Farris, 2010), implemented in the online code. Nevertheless, it was found to be more sensitive to noise than the proposed Feature Density measure. However, future work might consider it as an alternative measure of sparsity.

It is sometimes assumed that a single neural pattern underlies the observed decoding performance. But the brain is massively parallel, with cascades of overlapping neural events (Marti et al., 2015; Gwilliams & King, 2020; Desbordes et al., 2023; Hong et al., 2024). The methods described in this paper pave the way for more advanced tools that disentangle the multiple patterns in neural activity at any given time. Overall, we advocate for geometric reasoning, rather than a specific metric, as a powerful complement to decoding performance analyses.

The analysis of subspaces has emerged as a powerful way to gain insight into neural computation, for example in work showing that prefrontal populations separate working memory and motor preparation into distinct coding subspaces (Tang et al., 2020), as well as different subspaces for different items in working memory, thus reducing interference (Xie et al., 2022; Santo-Angles et al., 2025). Our method fits directly into this scheme by opening the possibility of testing subspace-based mechanisms in humans. Moreover, whereas orthogonal subspaces avoid interference, allowing encapsulated processing, a **progressive** rotation could reflect ongoing neural computation, analogous to mental rotation processes (Shepard & Metzler, 1971).

In summary, the Rotation Angle and Feature Density capture how the decision subspace of a linear decoder evolves in orientation and feature distribution over time. In doing so, these novel geometrical indexes complement classical decoding methods by not only indicating *when* information appears or generalizes, but also revealing *how* the underlying neural code evolves.

## 5. Acknowledgement

This research was funded by the National Center for Competence in Research Evolving Language (Swiss National Science Foundation Agreement #51NF40_180888, awarded to Nina Kazanina and Sophie Schwartz

