## Supplementary figures for "Temporal evolution of Neural Codes: The Added Value of a Geometric Approach to Linear Coefficients"

### Supplementary Materials

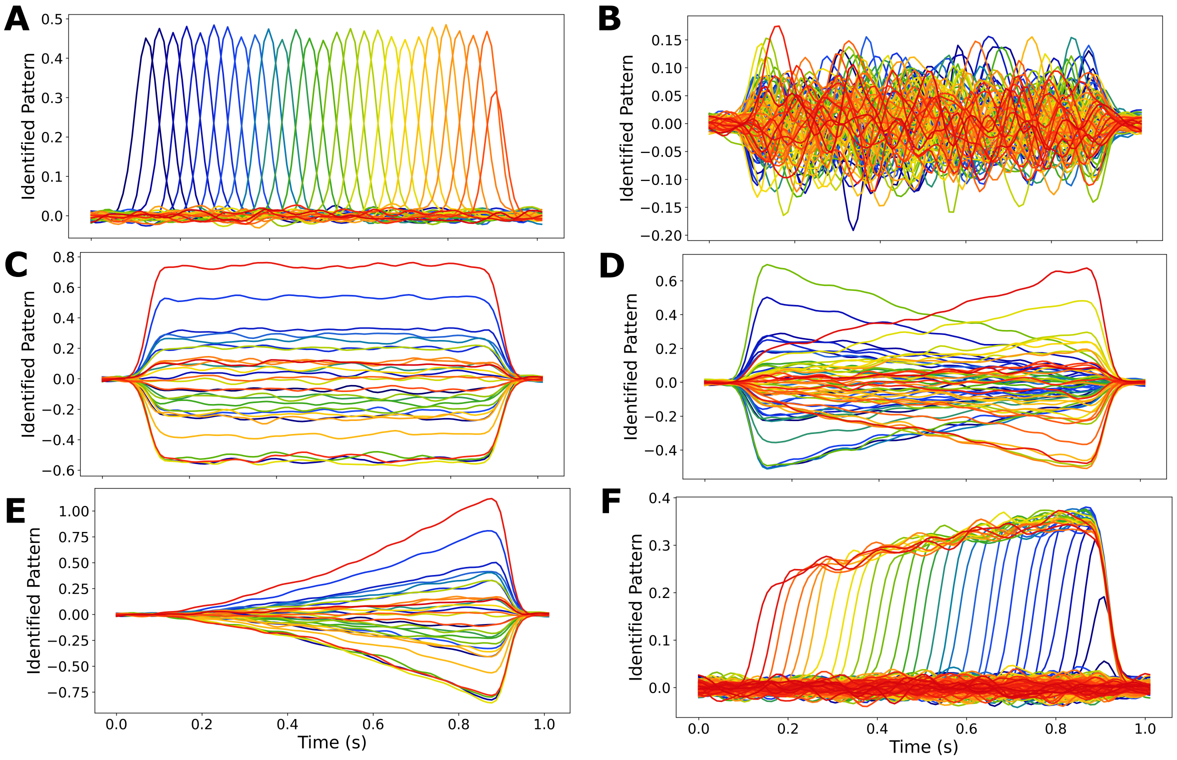


**Supplementary Figure 1: Identified patterns**

*Identified activity patterns for every scenario.*

***A**: Sparse Cascade, **B**: Distributed Cascade, **C**: Sustained, **D**: Progressive Switch, **E**: Scaling, **F**: Recruitment.*

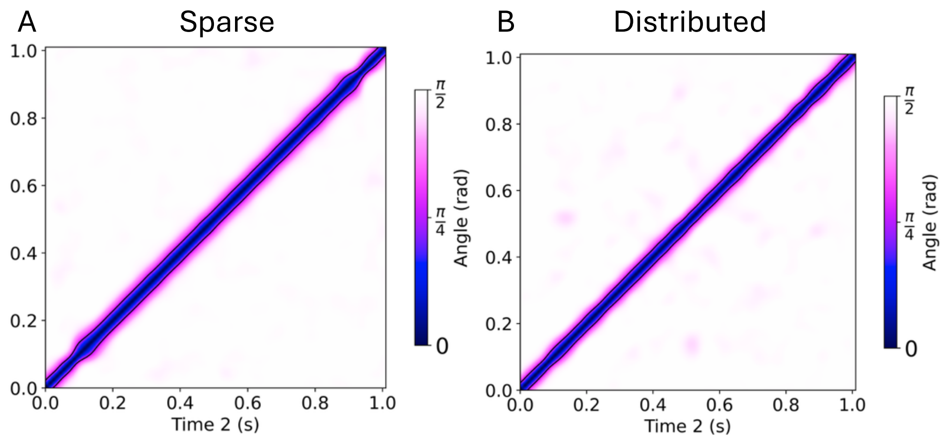


**Supplementary Figure 2: Rotation angle for the Sparse and Distributed scenarios**

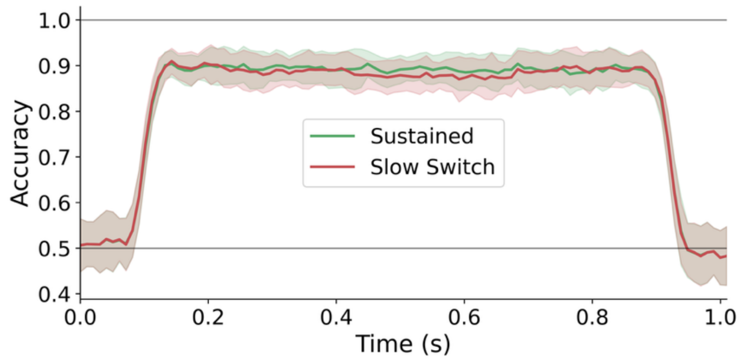


**Supplementary Figure 3: Within-time decoding performance for the Sustained and Progressive Switch scenarios**

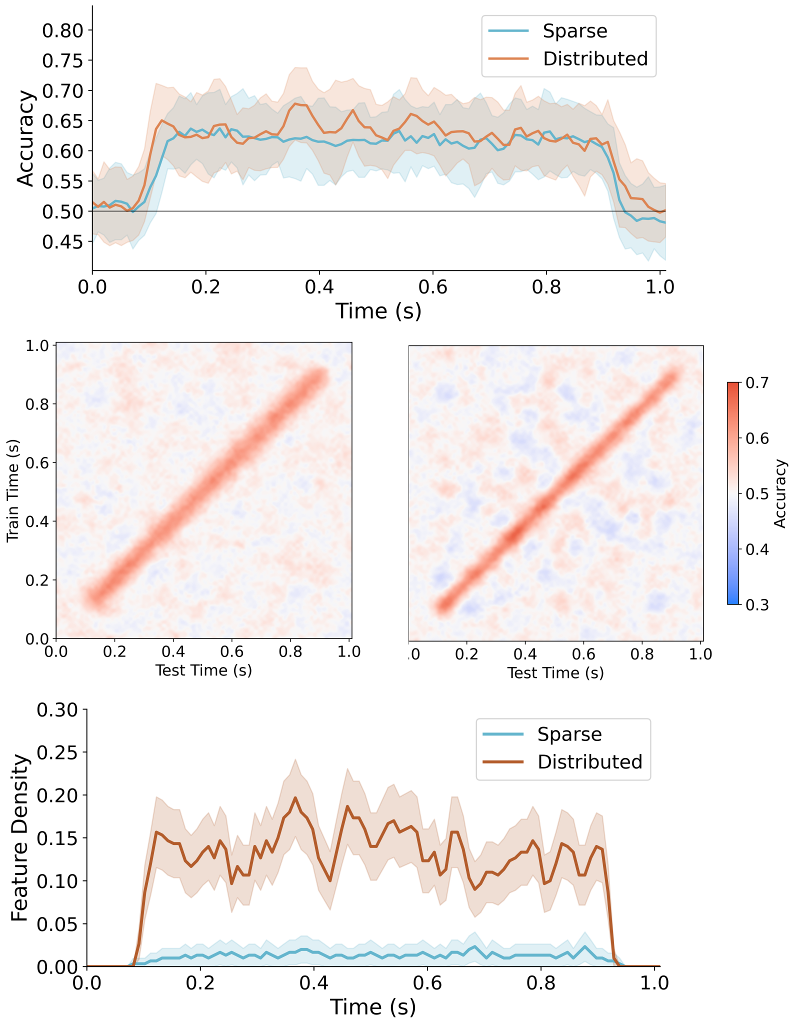


**Supplementary Figure 4: Classifier performance, generalization and Feature Density for Sparse and Distributed Sequential scenarios for Linear Support Vector Classifier**

***A**: Within-time decoding performance for both scenarios*

*Temporal generalization matrices for Sparse Cascade (**B**)and Distributed Cascade (**C**).*

***D**: Feature Density over time for both scenarios showing an index of the sparsity of the decision subspace.*

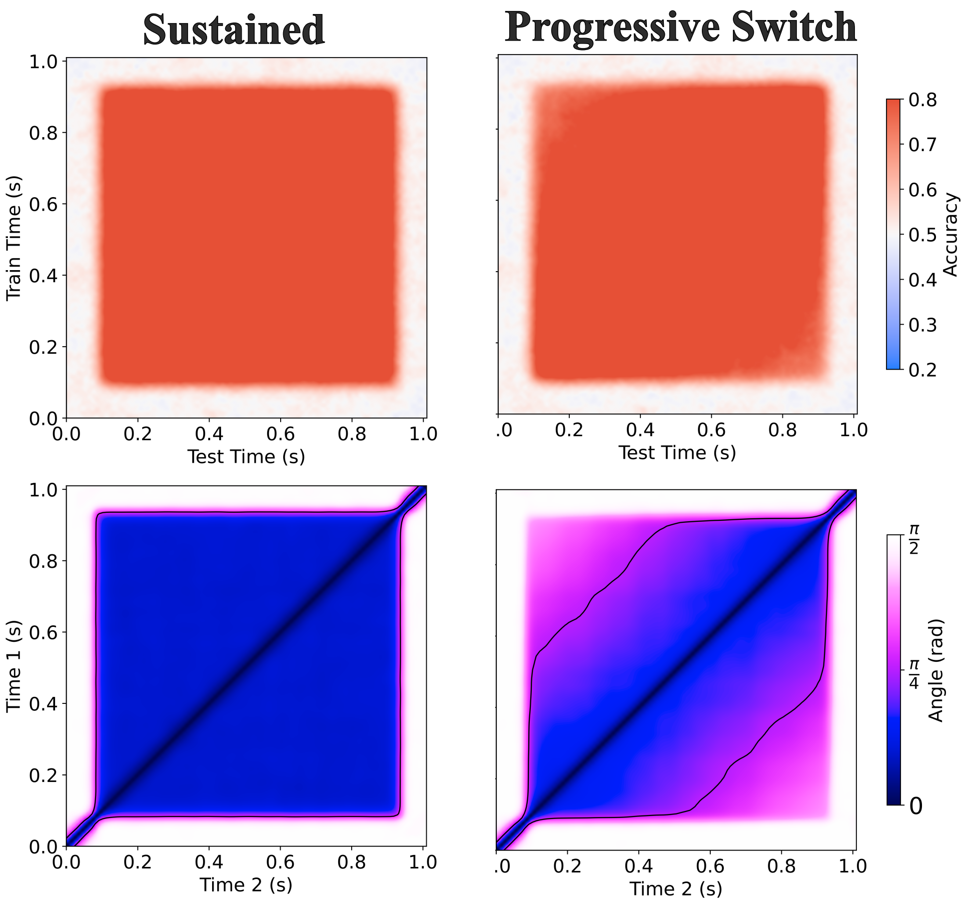


**Supplementary Figure 5: Decoding generalization performance and decision subspace Rotation Angle for Sustained and Progressive Switch scenarios with Linear Support-Vector Classifier**

*Temporal generalization matrices for Sustained (**A**)and Progressive Switch (**B**) scenarios.*

*Rotation Angle matrices for Sustained (**C**) and Progressive Switch (**D**). The thin black lines represent isoline of $\theta = 45$ °.*

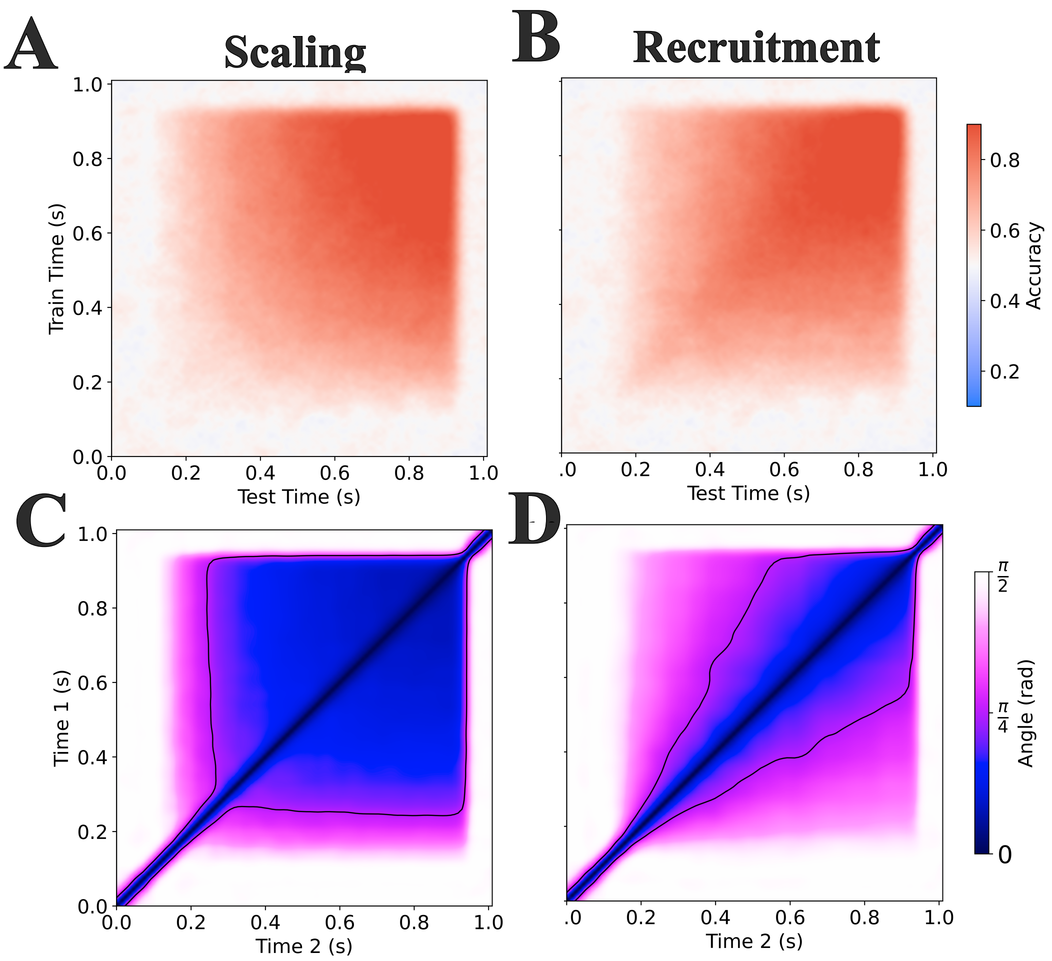


**Supplementary Figure 6: Decoding generalization performance and decision subspace Rotation Angle for Scaling and Recruitment scenarios with Linear Support-Vector Classifier**

*Temporal generalization matrices for Scaling (**A**)and Recruitment (**B**).*
*Rotation Angle matrices for Scaling (**C**) and Recruitment (**D**). The thin black lines represent isoline of* $\theta = \frac{\pi}{4} rad = 45°$. *Notice that for Scaling (C), after t = 0.3 s there is hardly any change in orientation of the decision subspaces (low angles), whereas in the case of Recruitment (D) for any row, there is an evolution of the colour witnessing the change in the angle.*

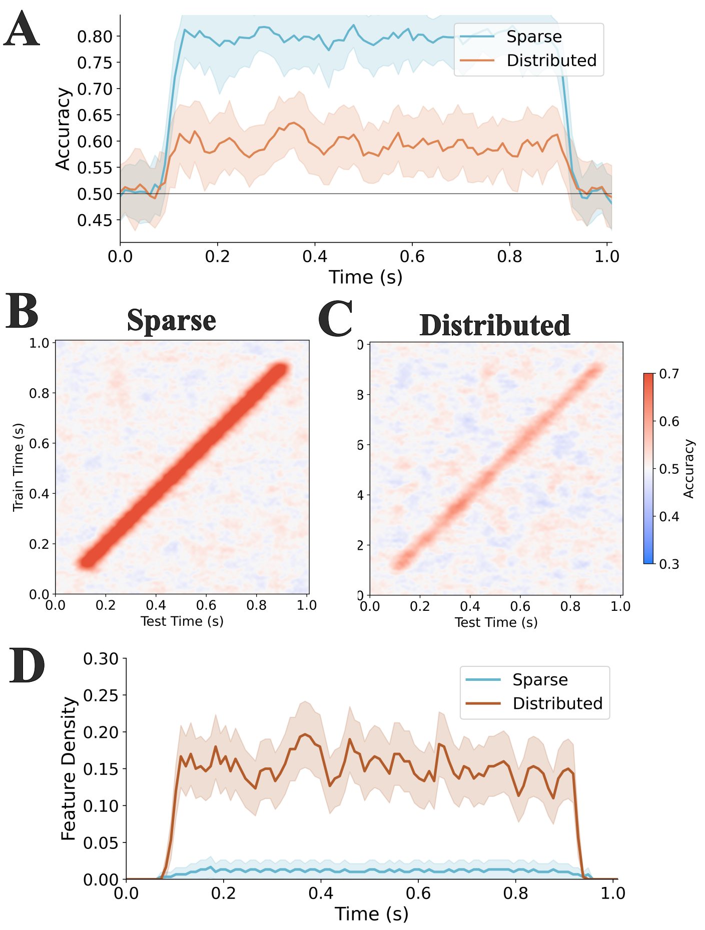


**Supplementary Figure 7: Classifier performance, generalization and Feature Density for Sparse and Distributed Sequential scenarios for Lasso regularized Logistic Regression**

***A****: Within-time decoding performance for both scenarios*

*Temporal generalization matrices for Sparse Cascade (****B****)and Distributed Cascade (****C****).*

***D****: Feature Density over time for both scenarios showing an index of the sparsity of the decision subspace.*

*Note that the Lasso regularization is better reaches better decoding performance on the sparse scenario. This is expected given that this methods finds a sparse solution.*

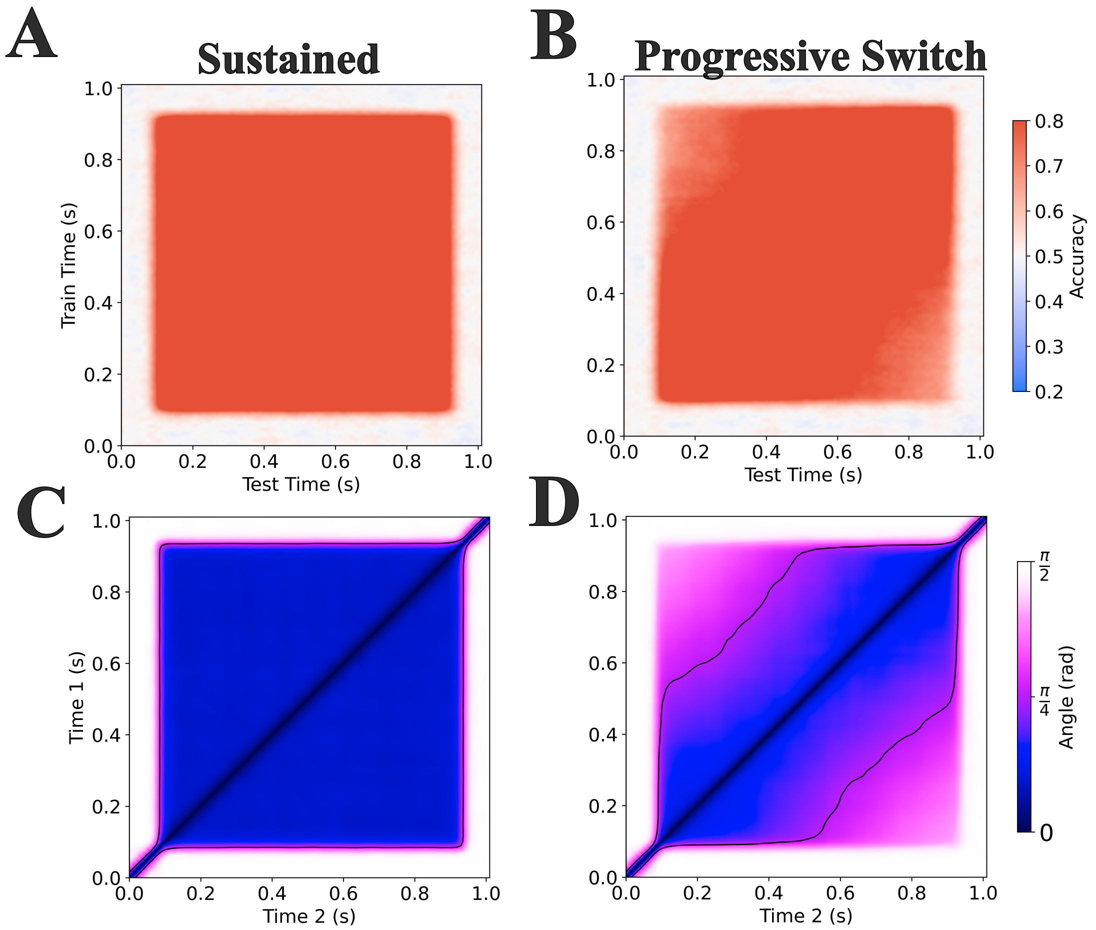


**Supplementary Figure 8: Decoding generalization performance and decision subspace Rotation Angle for Sustained and Progressive Switch scenarios with Lasso regularized Logistic Regression**

*Temporal generalization matrices for Sustained (**A**)and Progressive Switch (**B**) scenarios.*

*Rotation Angle matrices for Sustained (**C**) and Progressive Switch (**D**). The thin black lines represent isoline of* $\theta = 45°$.

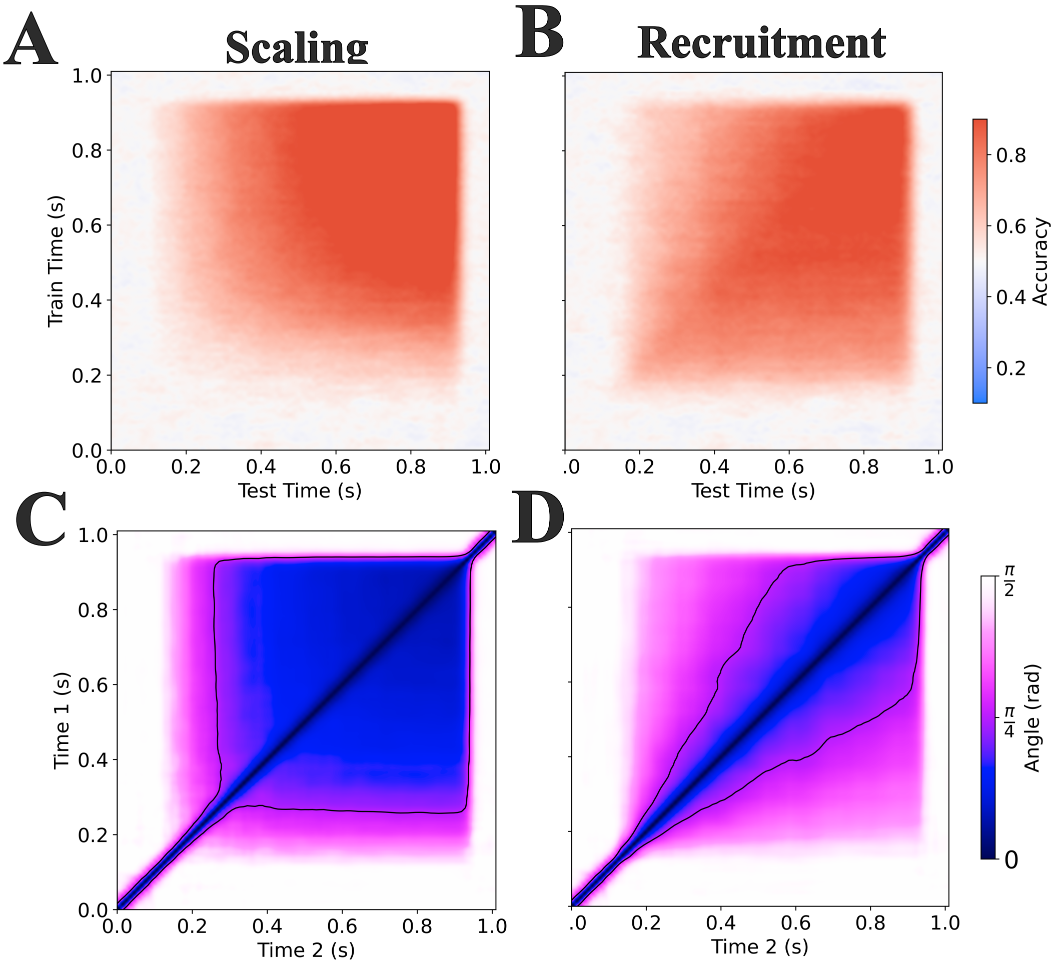


**Supplementary Figure 9: Decoding generalization performance and decision subspace Rotation Angle for Scaling and Recruitment scenarios with Lasso regularized Logistic Regression**

*Temporal generalization matrices for Scaling (**A**)and Recruitment (**B**).*
*Rotation Angle matrices for Scaling (**C**) and Recruitment (**D**). The thin black lines represent isoline of* $\theta = \frac{\pi}{4} rad = 45°$*. Notice that for Scaling (C), after t = 0.3 s there is hardly any change in orientation of the decision subspaces (low angles), whereas in the case of Recruitment (D) for any row, there is an evolution of the colour witnessing the change in the angle.*

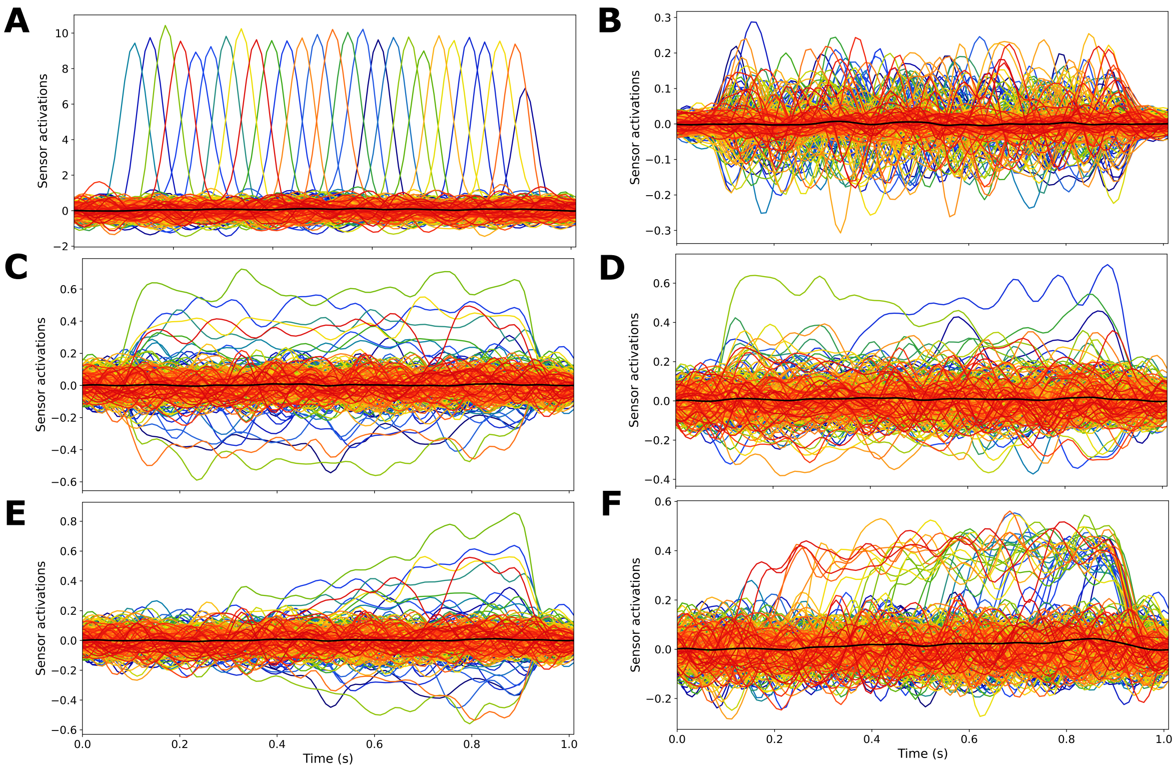


**Supplementary Figure 10: Evoked activations for each scenario**

*Average over all trials for each condition.*

***A**: Sparse Cascade, **B**: Distributed Cascade, **C**: Sustained, **D**: Progressive Switch,*
***E**: Scaling, **F**: Recruitment.*
